# Enhancing Intra-Continental Biogeographical Ancestry Prediction Through a Machine Learning Marker Selection Method

**DOI:** 10.1101/2025.11.08.687358

**Authors:** Theresa Maurer, Lennart Purucker, Frank Hutter, Peter Pfaffelhuber, Carola Sophia Heinzel

## Abstract

While classifiers such as TabPFN (Hollmann et al., 2025) and SNIPPER (Phillips et al., 2007a) achieve strong intercontinental performance (Heinzel et al., 2025), their accuracy in classifying individuals within Europe remains low. One major factor contributing to this limitation is the set of genetic markers used for classification. Marker panels such as the VISAGE Enhanced Tool (Xavier et al., 2022) are commonly employed in forensic genetics because they contain ancestry-informative markers (AIMs) that distinguish very well between major continental populations. However, these panels are often not optimized for fine-scale differentiation within continents, where genetic variation is more subtle and population structure is rather continuous.

We apply machine learning to select informative markers for intra-European classification, using data from Consortium et al. (2015). Compared with the VISAGE Enhanced Tool and allele frequency–based approaches (Phillips et al., 2007b; Kosoy et al., 2009; Nassir et al., 2009; Kidd et al., 2014; Phillips et al., 2014a), our marker sets achieve substantially higher accuracy within Europe: For four European populations, accuracy improves from 68.2% (VISAGE, 104 markers) to 73.7% (100 new markers) and 82.3% (200 new markers). For five populations, accuracy rises from 56.1% (VISAGE) to 64.5% (100 new markers).

Our results show that tailored marker selection markedly improves intra-continental classification. While optimized here for Europe, the method can be applied to any region with sufficient training data.

## 1 Introduction

Classification of individuals into ancestral populations, i.e. inferring biogeographical ancestry (BGA) from DNA traces, is a key task in forensic science (Wen et al., 2023; Phillips et al., 2009, 2007b; Dang et al., 2005; Min et al., 2000; Gan et al., 2003). Individuals are commonly classified into continental groups such as Africa, Europe, East Asia, South Asia, Admixed Americans, and Oceania (Cheung et al., 2017; Wang et al., 2007; Mogensen et al., 2020), but finer-scale intra-continental classification is also of growing interest. The performance of a classifier is assessed using measures such as misclassification rate, logloss, and ROC AUC. Population classification relies on two steps: choosing the best classifier and choosing the best marker set for this classifier and for the given data. Heinzel et al. (2025) showed that the machine learning method TabPFN (Hollmann et al., 2025) performs best for the classification into ancestral populations. The VISAGE Enhanced Tool, combined with the TabPFN classifier, has shown high accuracy for inter-continental classification, but leaves room for improvement at the intra-continental level. This motivates our focus on improving marker selection specifically for intra-continental classification, where it has a major impact on performance (Cheung et al., 2017).

Traditionally, Ancestry Informative Markers (AIMs) are chosen based on large allele frequency differences between populations (Phillips et al., 2007b; Kosoy et al., 2009; Nassir et al., 2009; Phillips et al., 2014a; Ruiz-Ramírez et al., 2023). Various other strategies for marker selection have been explored. For example, openADMIXTURE provides an unsupervised approach that avoids labeled training data (Ko et al., 2023). Multi-InDels have been proposed as alternatives to AIMs (Sun et al., 2022, 2019). Other approaches include informativeness measures (Phillips et al., 2014a), *F*_*ST*_ -based criteria (Kidd et al., 2014), stepwise feature selection (Pfaffelhuber et al., 2020b), or admixture model–based methods (Pfaff et al., 2004; Alexander et al., 2009; Pfaffelhuber and Rohde, 2022; Heinzel, 2025). Practical panels, such as the VERO-GEN Enhanced Tool (Xavier et al., 2022), combine different strategies, while some approaches also integrate phenotypic traits (Bulbul et al., 2016) or multi-stage filtering (Toma et al., 2018). Pilli et al. (2023) combined markers from previous marker sets.

However, the existing marker selection methods are not tailored to the classification task they are later applied to. In particular, it remains unclear whether criteria such as allele frequency differences are truly optimal for classification.

In this work, we address this gap by applying machine learning methods to directly select markers optimized for the classification task at hand. The goal is to develop a method that identifies small, highly informative marker sets (50–200 markers) instead of thousands (Ko et al., 2023), thus reducing sequencing effort.

## 2 Material and Methods

Figure 1 visualizes our approach. Each step that is mentioned in this figure is described in more detail below.

**Figure 1.**
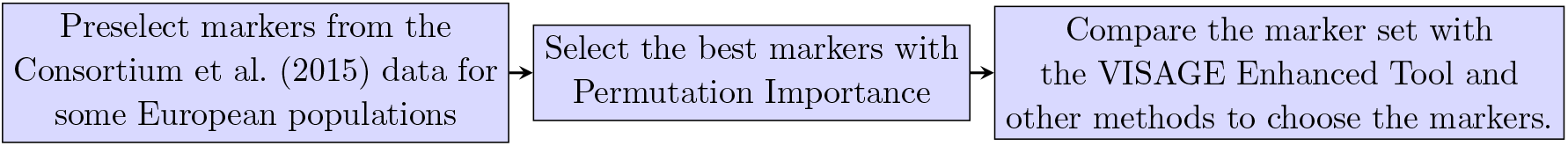
Overview of the marker selection and the evaluation of the methods.

### 2.1 Choice of the Training Data

We use four (Finland (FIN), Toscani in Italy (TSI), Iberian Population in Spain (IBS) and British in England and Scotland (GBR)) out of fives European populations within Consortium et al. (2015) to select the markers.

Based on these individuals, we preselect markers that have a minor allele frequency of at least 5% in the populations in total. After that, we only use every 2.000^*th*^ allele. This has two reasons: many markers that are closely together on a chromosome usually do not provide more information than one of these markers due to linkage and the computational cost increases exponentially in the number of markers. Thereby, we get approximately 6000 bi-allelic markers. In addition to that, we also take the 104 markers from the VISAGE Enhanced Tool into account.

### 2.2 Selection of the Markers

Our goal is to select a subset of the ∼6000 markers such that we maximize the predictive performance of machine learning methods. In other words, we aim to solve a feature selection problem (Saeys et al., 2007; Tang et al., 2014; Venkatesh and Anuradha, 2019) for our static high-dimensional tabular data (Li et al., 2017).

While studying benchmarks of feature selection for high-dimensional data, we were unable to find conclusive evidence recommending a single best feature selection method for state-of-the-art deep learning models, such as TabPFN, for high-dimensional data (cf. (Bommert et al., 2020; Hopf and Reifenrath, 2021; Li et al., 2022; Cherepanova et al., 2023)). We hypothesize that this is due to the lack of using strong and well-tuned deep learning models in prior benchmarks. Even more recent work, such as that by Cherepanova et al. (2023), is already outdated because of recent progress in deep learning for tabular data. Thus, we began exploring various feature selection methods, which have yielded the best performance in prior benchmarks, through preliminary experiments. However, most of the methods we tried did not yield competitive results in early trials. Their implementations were either unable to scale to our ∼6000 markers, or selected features resulted in poor performance for any machine learning models we tested. In particular, we observed that tree-based feature selection methods were unable to produce well-performing features for deep learning methods.

At the same time, we noticed that methods based on permutation importance (Janitza et al., 2018; Wright et al., 2016; Parr et al., 2018) performed the best among all the other choices. This aligned with the competitive performance of the permutation importance implementation based on random forests from ranger (Wright and Ziegler, 2017) in prior benchmarks. Yet, to the best of our knowledge, no prior work has evaluated feature selection with state-of-the-art deep learning methods using permutation importance. By design, permutation importance is model-agnostic and is therefore used in methods such as SHAP (Lundberg and Lee, 2017) as part of the model-agnostic explainer, or by popular predictive machine learning systems, such as AutoGluon (Erickson et al., 2020), as the default method for computing feature importance. Thus, we chose to employ permutation importance with a deep learning model (PI-DL) to determine a subset of the ∼6000 markers.

### Permutation Importance

Permutation importance of a feature measures how much model performance drops when a feature’s values are randomly shuffled. A score of 0.01 indicates a 0.01% decrease in performance. Higher scores indicate greater importance, while negative scores suggest the feature may harm performance and that removing it could improve results.

### A Deep Learning Model for Permutation Importance

Permutation importance can be computed with any model where single features can be shuffled and performance can be measured with little cost. Thus, to select the best deep learning model, we surveyed benchmarks for tabular data and selected RealMLP (Holzmüller et al., 2024), TabM (Gorishniy et al., 2025), and TabPFN (Hollmann et al., 2025) as candidates. These candidates were the best performing deep learning methods in the recent TabArena benchmark (Erickson et al., 2025). TabPFN is limited to 500 features due to pretraining, so we ruled it out, as we did not expect its performance to generalize to 6000 features, which would make its permutation importance misleading. In preliminary experiments, RealMLP performed worse than TabM when being trained on all 6000 features. Thus, we ultimately selected TabM.

TabM is a recently introduced, lightweight deep learning model based on a multilayer perceptron architecture. Due to its lightweight implementation and optimization for GPU usage, we were able to train a model on our high-dimensional data faster than with any other model we tested in preliminary experiments, while maintaining very high predictive accuracy when using all features.

### The Marker Selection Pipeline

To employ permutation importance with a deep learning model for marker selection, we opted for the following pipeline, which we also share on our public GitHub repository^1^: (1) Given a training data set, we used TabM to compute permutation importance with five shuffle sets and 4-fold inner cross-validation, using the implementation from AutoGluon (Erickson et al., 2020). Therefore, we trained TabM and performed a 4-fold inner cross-validation in parallel on one NVIDIA L40S with 48 GB of VRAM, utilizing up to 1 hour of training time. (2) We repeat the first step 60 times, by performing 20-repeated 3-fold outer cross-validation. In other words, we split our original data 60 times into train-test sets to guard against randomness in the training data before computing permutation importance. In the end, we obtain the feature importance for all ∼6000 markers 60 times. We parallelized these repetitions across several GPUs to reduce the higher one-time cost of this exhaustive process. (3) Finally, to produce one subset of markers we can suggest to practitioners, we average the permutation importance over all repeats and take the top 50, 100, or 200 features with the highest permutation importance. Some extended information about the top 100 markers can be found in section 7.4. Note that our marker set might be slightly overfit to the available data, as we use the available data to determine the marker set.

### 2.3 Evaluation of the Marker Sets

We evaluate each marker set by training the model, either TabPFN on a subset of the data and evaluating its prediction on a holdout set, i.e. we use Cross Validation. We compare the performance of PI-DL with the best 50, 100, 200 markers with the VISAGE Enhanced Tool, the preselected markers set that consists of about 6000 markers and using the differences between the allele frequency as e.g. suggested by Kidd et al. (2014); Phillips et al. (2014b); Kosoy et al. (2009). More precisely, for the last method, we calculate for each marker and for each pair of populations the absolute difference between the allele frequencies. An overview of the different cases that will show up can be found in Table 1.

**Table 1:**
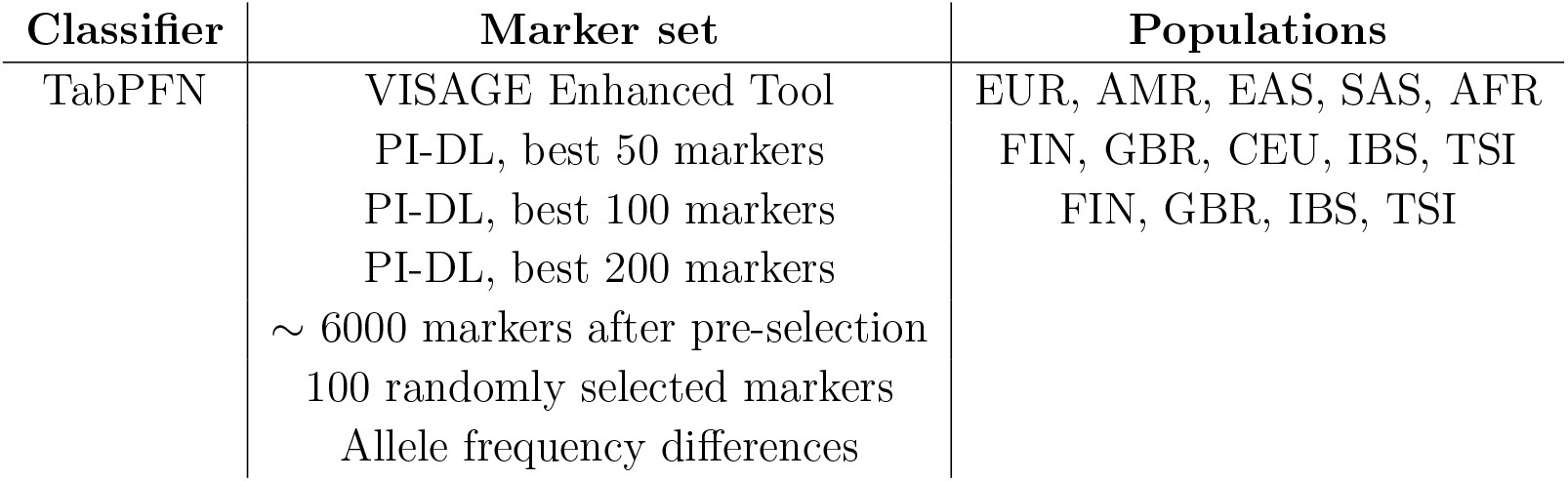
Overview of the different cases that we consider here to compare the marker sets to each other. We considered all possible combinations of the entries in the columns. The 100 randomly selected markers refer to 100 randomly selected markers from the ∼ 6000 markers that we get after the pre-selection.

Let us consider the selection based on allele frequency differences in more detail. Therefore, let *K* be the number of populations. For each bi-allelic marker, there are (*K* − 1)!(= 6 in case *K* = 4)possible pairwise allele frequency differences between populations. For each of these six population pairs, we rank all markers by the absolute allele frequency difference. We then select the top ⌊100*/*6⌋ = 16 markers with the highest differences for each pair. This procedure does not result in exactly 6 *×* 16 = 96 distinct markers, as some markers may appear among the top-ranked ones for multiple population pairs. To reach a total of 100 unique markers, we continue selecting the next-best markers from each pairwise ranking—excluding those already chosen—until the full marker set contains 100 unique entries.

We compare the performance of the marker sets not only for the populations for which the marker set is trained, but also for unseen populations. More precisely, we compare the different marker sets for (i) FIN, TSI, IBS, GBR (4 populations), (ii) FIN, TSI, IBS, GBR and from Utah Residents (CEPH) with Northern and Western European Ancestry (CEU) (5 populations) and (iii) Africa (AFR), Europe (EUR), South Asia (SAS), East Asia (EAS), Admixed Americans (AMR) (5 populations). The percentage of individuals from the populations within the data set is shown in Figure 2, while the number of individuals per population can be found in Table 2.

**Table 2:**
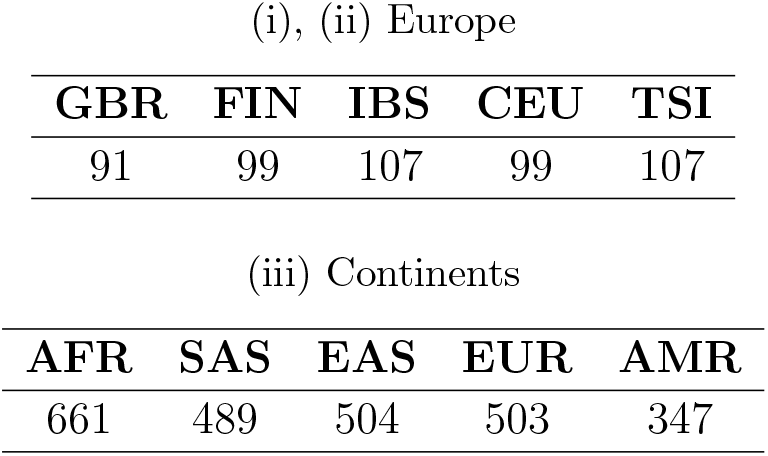
Numbers of individuals per population for case (i), (ii) and (iii).

| GBR | FIN | IBS | CEU | TSI |
| --- | --- | --- | --- | --- |
| 91 | 99 | 107 | 99 | 107 |

Table 2: Numbers of individuals per population for case (i), (ii) and (iii).
| AFR | SAS | EAS | EUR | AMR |
| --- | --- | --- | --- | --- |
| 661 | 489 | 504 | 503 | 347 |

**Figure 2.**
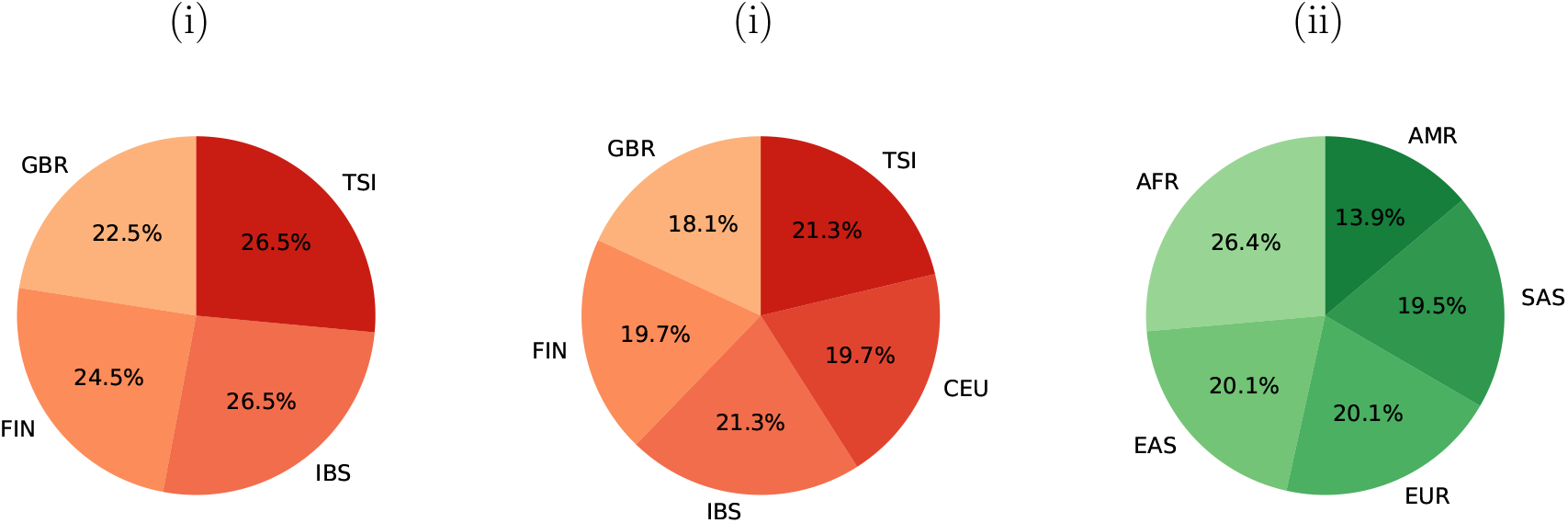
Percentage of individuals per population for the cases (i), (ii) and (iii). For exact numbers, see Tables 2.

This experiment is repeated 50 times by using 10-repeated 5-fold cross-validation with stratification from scikit-learn (Pedregosa et al., 2011), respectively. We score predictions using accuracy, log loss and ROC AUC^2^, following best practices from machine learning (Provost et al., 1998; McElfresh et al., 2023; Gijsbers et al., 2024; Vovk, 2015). ROC AUC measures the ability to distinguish between classes, where 1, indicating perfect discrimination, and 0.5, representing random guessing. Analogously to previous evaluation metrics in forensics (Pfaffelhuber et al., 2020a; Resutik et al., 2023; Heinzel et al., 2025), we also count the number of true positive and false negative samples to compute a confusion matrix. More precisely, for every of our 50 or 60 repetitions, we sum up the number of individuals from class *i*, e.g., Spain, that are classified into population *j*, e.g., Italy. The resulting percentiles are depicted as a confusion matrix.

### 2.4 Code availability

The code for our analysis can be downloaded from GitHub. On this website you can also find an overview of the hyperparameter that we used and the versions of the Python packages. We include mostly Python scripts for selecting individuals from the dataset, and a locally running web-interface for easier use of classification. We also set up an interface online and offline.

## 3 Results

### 3.1 Metrics

The ROC AUC (between 0.5 and 1, higher is better), Accuracy (between 0 and 1, higher is better) and the Log loss (greater than 0, lower is better) of the classification in case (i) can be found in Figure 3 (the numerical values for GBR, IBS, TSI, and FIN can be found in Section 7.3).

**Figure 3.**
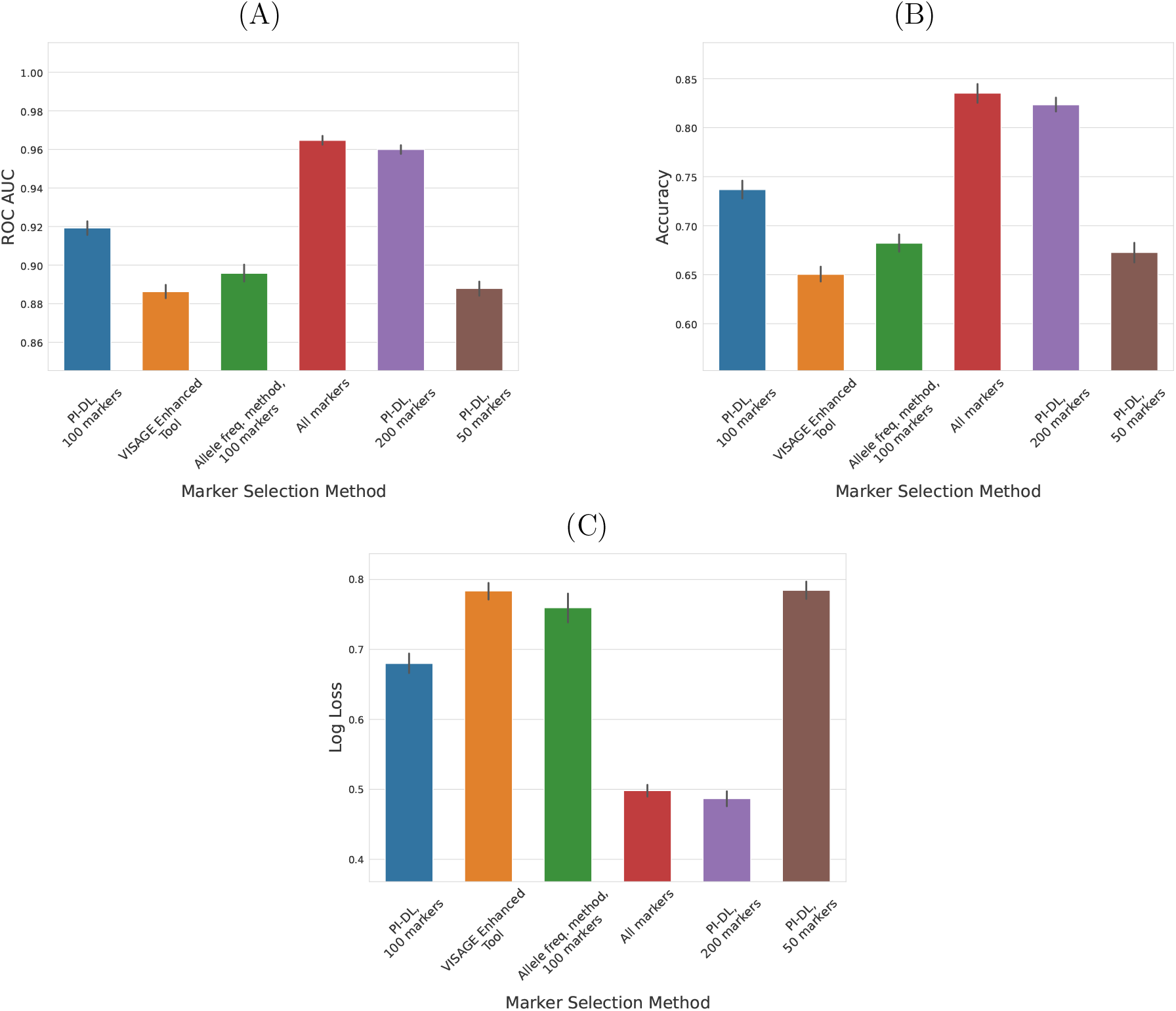
Evaluation metrics for inter-continental classification with TabPFN. The black line represents the 95% confidence interval. (A) ROC AUC; (B) Accuracy; (C) Log Loss.

For ROC AUC and Accuracy, the whole data set with about 6000 markers performs the best, while our novel marker set with 200 markers is the best regarding the Log Loss. However, of course, the comparison of markers sets with different number of markers is unfair towards the marker sets with the fewer markers. If we compare our novel marker set with 100 markers to the one by Xavier et al. (2022) that consists of 104 markers and the one that is used by using the maximal allele frequency difference, we still see that our novel method outperforms the previous marker selection methods. Note that TabPFN was only tested with up to 500 features. Running it with about 6000 markers requires a high computational power.

Altogether, our novel marker set that consists of either 100 or 200 markers, and the data set with about 6000 markers perform well, even for the intra-continental classification. We also see that the marker set with 50 markers that was created by using our novel method slightly better than the marker set by Xavier et al. (2022). Here, again, the comparison is unfair since the marker set by Xavier et al. (2022) is not optimized with respect to these specific populations. The results for case (ii) are shown in Figure 4. Again, we see that both the ROC AUC and the Accuracy of the newly chosen marker set with 100 features are much higher than of the VISAGE Enhanced Tool. Furthermore, the figure shows that the Log Loss of the markers that have been selected by using PI-DL, is much lower than for the VISAGE Enhanced Tool. This is remarkable, since the data contains one population, CEU, that has not been used to select the markers.

**Figure 4.**
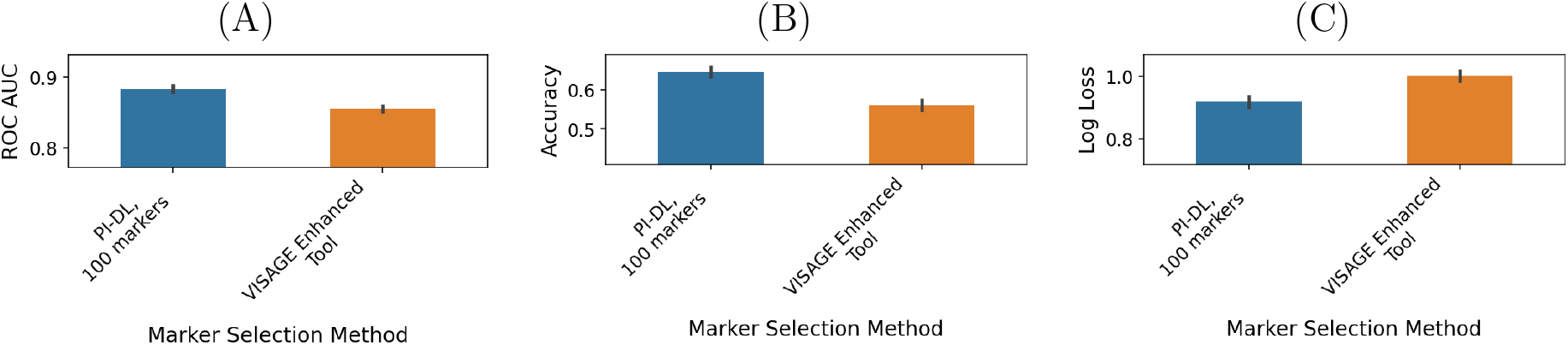
Evaluation metrics for intra-continental classification with TabPFN with CEU, TSI, IBS, GBR and FIN as populations. The black line represents the 95% confidence interval. (A) ROC AUC; (B) Accuracy; (C) Log Loss.

Figure 5 shows the metrics for the classification at an inter-continental level, i.e. for case (iii). This figure shows that the classification at a continental level works very well with both marker sets, the VISAGE Enhanced Tool and the 100 markers that have been selected with PI-DL.

**Figure 5.**
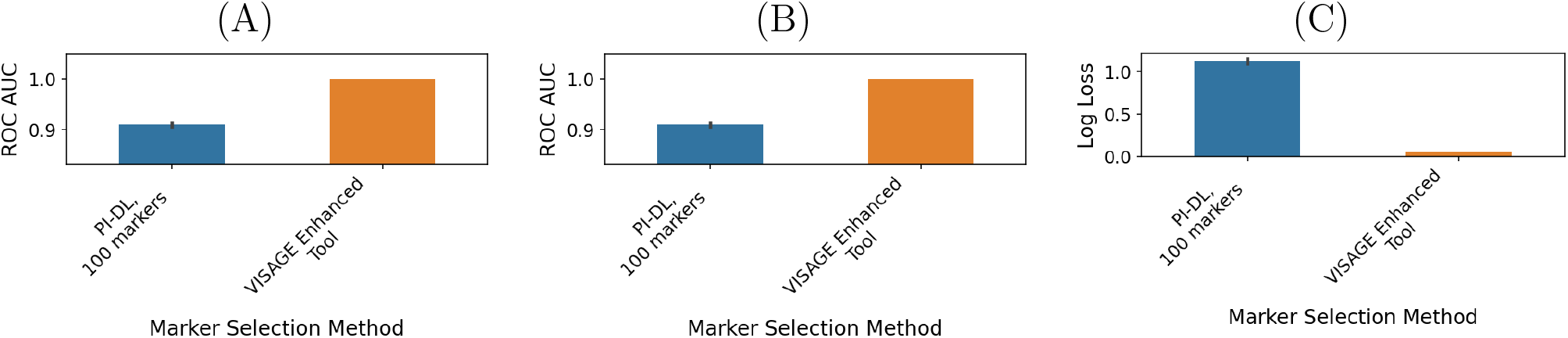
Evaluation metrics for inter-continental classification with TabPFN with EAS, SAS, AMR, AFR and EUR as populations. The black line represents the 95% confidence interval. ROC AUC; (B) Accuracy; (C) Log Loss.

Here, we clearly see that the VISAGE Enhanced Tool performs better than the 100 markers that has been chosen with PI-DL. The reason for this is that we did not train PI-DL to distinguish between continental populations.

### 3.2 Confusion Matrices

For more precise results which distinctions work well and which don’t, we display confusion matrices for all seven marker sets.

Figure 6 shows the confusion matrices for TabPFN of our novel marker set and the VIS-AGE Enhanced marker set for (ii) five European populations and (iii) the inter-continental classification.

**Figure 6.**
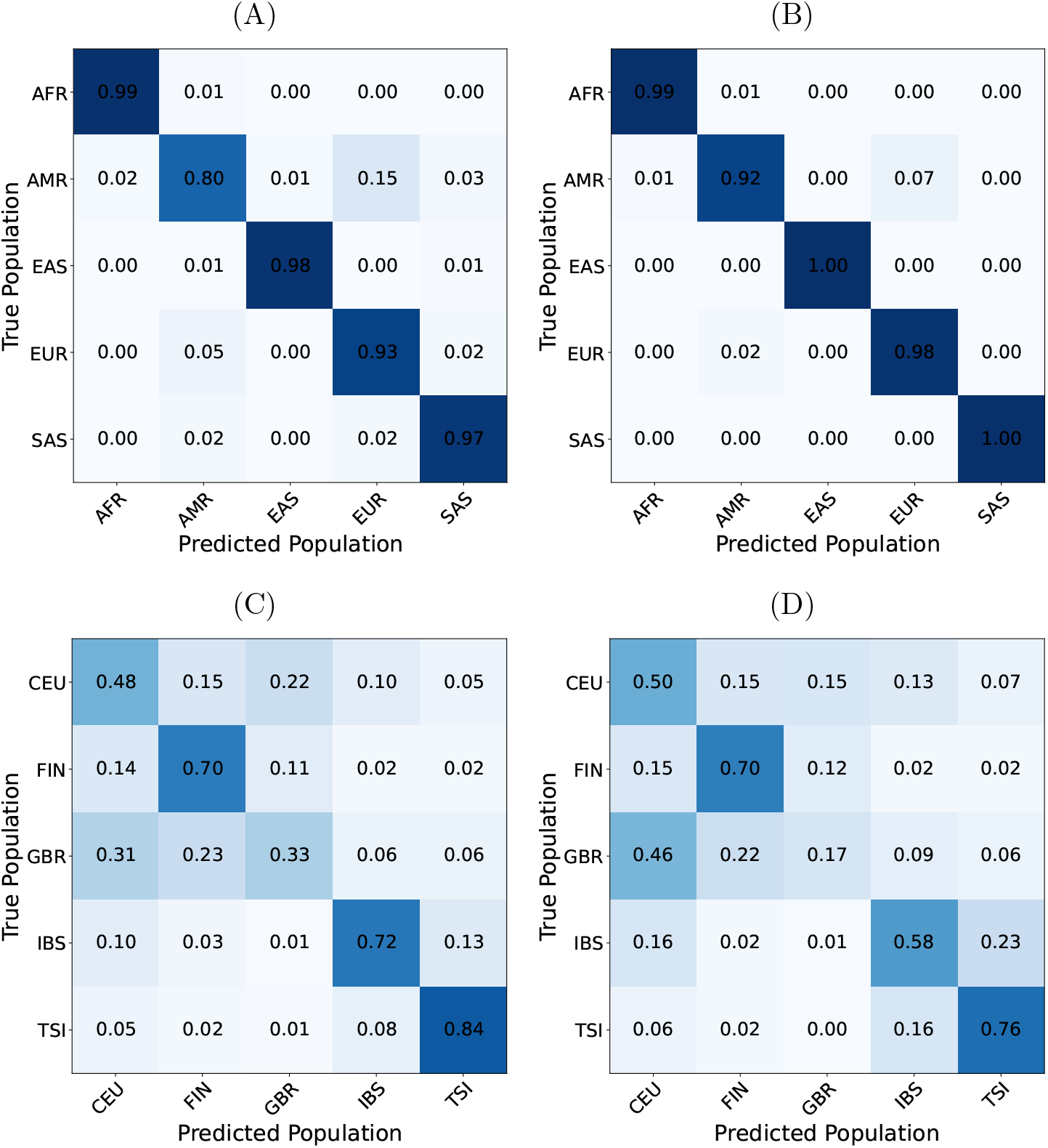
Confusion matrices for classification with TabPFN. All rows sum to 1 up to a rounding error. (A) Continental Classification, top 100 PI-DL, (B) Continental Classification, VISAGE Enhanced Tool, (C) Intra-Continental Classification, top 100 PI-DL and (D) Intra-Continental Classification, VISAGE Enhanced Tool

For the inter-continental classification, these confusion matrices show that both marker sets perform comparably well for the classification into AFR, EAS and SAS. However, the percentage of correct classified individuals from Europe is with the new marker set only 93% compared to 98% with the VISAGE Enhanced Tool. The biggest difference occurs for AMR: Here, the percentage of correct classified individuals from AMR with the new marker set is much lower than with the VISAGE Enhanced Tool. These results show that the classification between AFR, EAS, EUR and SAS works, even if the training data is only from Europe. If researchers also want to differentiation between AMR and EUR, some new markers have to be added to this marker set.

For the classification of individuals to CEU, FIN, GBR, IBS and TSI, we see that the new marker set works much better for IBS and TSI than the VISAGE Enhanced Tool. We also see that the confusion between GBR and CEU is high for both marker sets. However, the novel marker set can differentiate better between these two populations even though we did not train them on CEU.

## 4 Discussion

We compared different marker sets for intra-European classification. Our results show that the proposed method achieves the highest average ROC AUC, the highest average accuracy, and the best log loss when comparing marker sets of approximately 100 markers. For example, using TabPFN with our marker set with 100 markers improves accuracy from 56.1% with the VISAGE panel Xavier et al. (2022) to 64.5%.

Based on these results, we recommend our method for marker selection. Of course, before performing any marker selection, the required classification task (e.g. inter- or intra-continental, or both), must be fixed. To this purpose, we provide an offline interface with example data for simple application of our method to new data, e.g. for intra-African classification. In practice, we also advocate using our proposed marker set for the intra-European classification.

Interestingly, classification performance shows little difference between using 200 and 6,000 markers. However, with advances in sequencing technology (Satam et al., 2023; Kumar et al., 2024), using larger sets—for example, 200 instead of 104 markers as in the VISAGE panel—has become increasingly feasible.

Further improvements may also be achieved by refining the pre-selection of markers. Therefore, researchers could e.g. use openADMIXTURE (Ko et al., 2023) that uses a sparse *K*-means with feature ranking method to identify a subset of a number of markers that is small enough to serve as a input data set of our algorithm Ideally, selection should be performed from the complete set of available markers rather than from a restricted subset. Alternatively, the pre-selection could also be optimized by using only markers for which is already known that they provide information about the ancestry. This would be the same approach as used by Armstrong et al. (2025).

Finally, the optimal approach would jointly identify both the best classifier and the best marker set. In this work, however, we applied a two-step strategy: first identifying the best classifier for a given marker set Heinzel et al. (2025), and then determining the optimal marker set for that classifier.

## 7 Appendix

### 7.1 Number of Individual per Population

Table 2 shows the number of individuals per population.

### 7.2 Confusion Matrices

We present the confusion matrices for 100 random markers, PI-DL with 50 markers and all markers in Figure 7.

**Figure 7.**
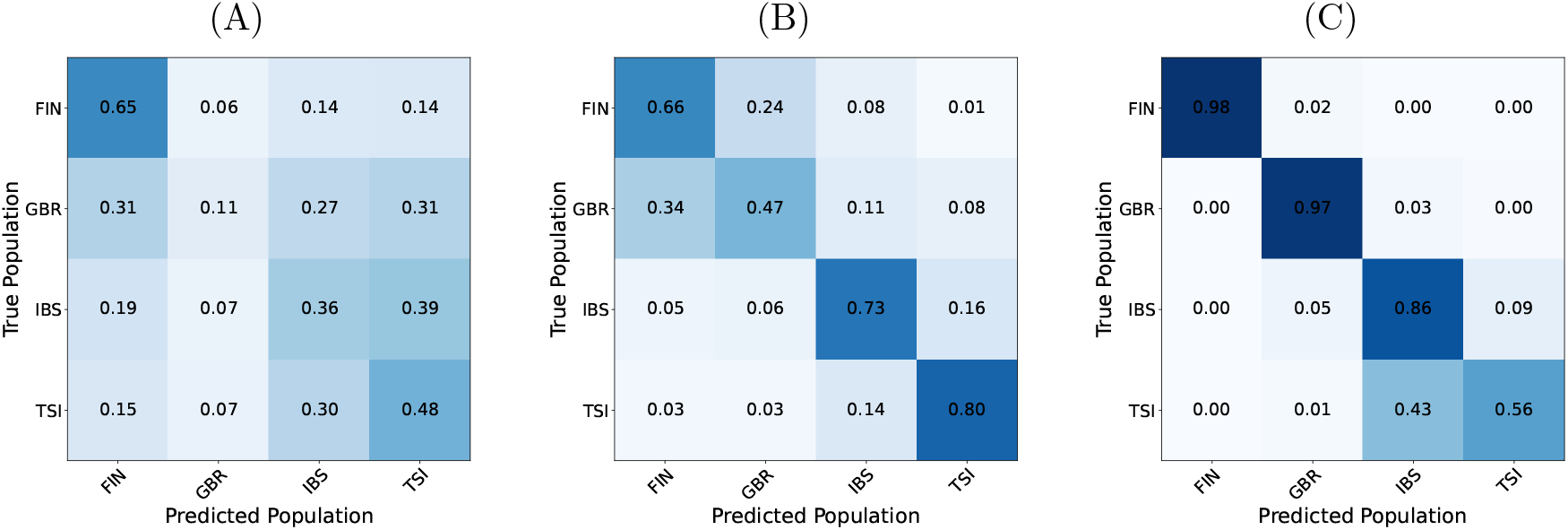
Confusion Matrices intra-continental classification with TabPFN. (A) 100 random markers, (B) PI-DL, 50 markers, (C) all markers.

### 7.3 Absolute Values Metrics

Table 3 shows the absolute values of the metrics Accuracy, ROC AUC, Balanced Accuracy and Log Loss for the populations IBS, TSI, GBR and FIN and for TabPFN.

**Table 3:** Evaluation Metrics for TabPFN.

| Feature Selection Method | Accuracy | Balanced Accuracy | ROC AUC | Log Loss |
| --- | --- | --- | --- | --- |
| New Method (100 Features) | 0.736781 | 0.731706 | 0.919263 | 0.679918 |
| Expert knowledge (104 Features) | 0.650379 | 0.647380 | 0.886219 | 0.783562 |
| Allele Freq. Method (100 Features) | 0.682146 | 0.684132 | 0.895750 | 0.759571 |
| 100 Random Selected Features | 0.408149 | 0.401769 | 0.670939 | 1.257144 |
| All features | 0.835173 | 0.842948 | 0.964697 | 0.498137 |
| New Method (200 Features) | 0.823274 | 0.821199 | 0.959955 | 0.486721 |
| New Method (50 Features) | 0.672677 | 0.664847 | 0.887899 | 0.784496 |

### 7.4 Extended Information about the marker set

Since e.g. the Admixture Model assumes that the loci are unlinked, we also present the genetic distance between the loci in Figure 8.

**Figure 8.**
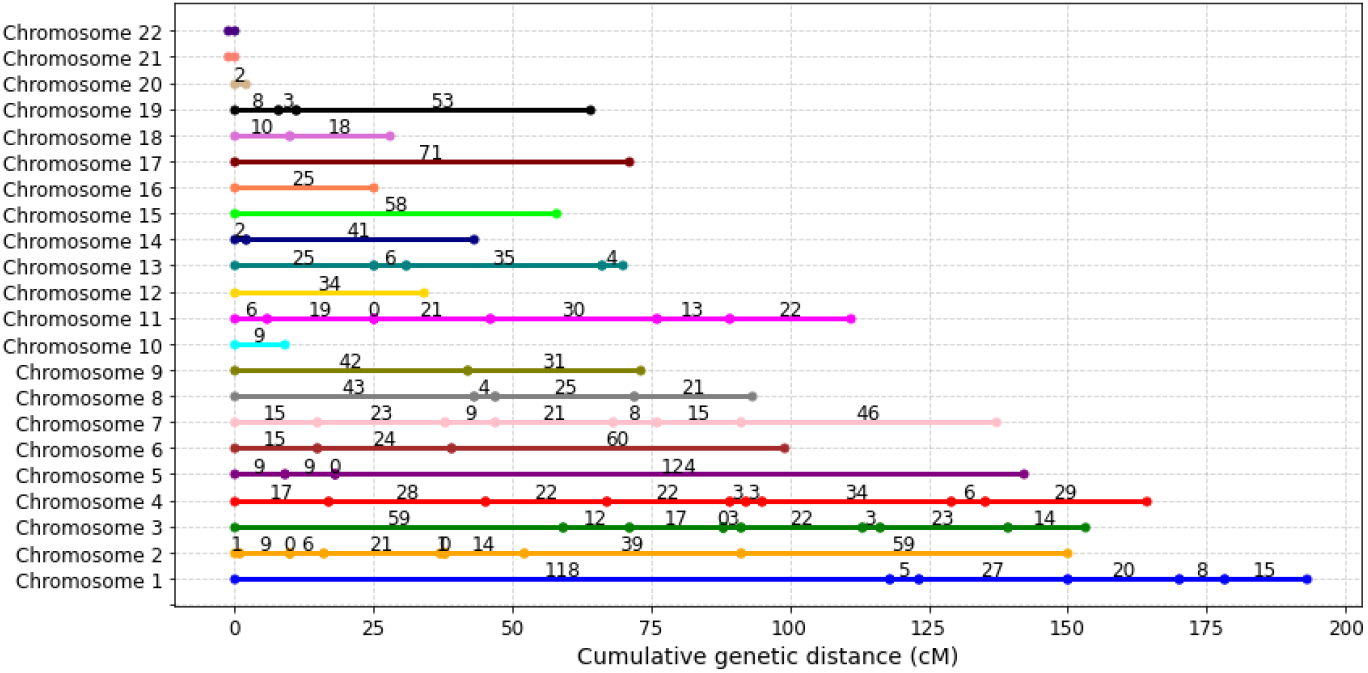
Distance between the markers of our marker set in centi Morgan (cM).

Table 4 shows the position of the 100 markers that we get by using PI-DL.

**Table 4:** The 100 best markers with our method. They are ordered according to their position on the chromosome. Blue indicates that these markers are also included in the VISAGE Enhanced marker set.

| Chrom | Position | Chrom | Position | Chrom | Position | Chrom | Position |
| --- | --- | --- | --- | --- | --- | --- | --- |
| 1 | 10177 | 1 | 89531508 | 1 | 94539004 | 1 | 120222104 |
| 1 | 158785947 | 1 | 164392446 | 1 | 177206543 | 2 | 89342505 |
| 2 | 89830663 | 2 | 109359940 | 2 | 109513601 | 2 | 115140035 |
| 2 | 135564058 | 2 | 136707982 | 2 | 136965117 | 2 | 153795774 |
| 2 | 202254288 | 2 | 242300443 | 3 | 7265212 | 3 | 60125368 |
| 3 | 67131402 | 3 | 80348904 | 3 | 82952542 | 3 | 95908174 |
| 3 | 121459589 | 3 | 123942108 | 3 | 147771903 | 3 | 165792726 |
| 4 | 19635773 | 4 | 31847585 | 4 | 65877053 | 4 | 90115375 |
| 4 | 116345962 | 4 | 120750919 | 4 | 124354293 | 4 | 162394576 |
| 4 | 167399504 | 4 | 185618037 | 5 | 28037143 | 5 | 45455862 |
| 5 | 49405663 | 5 | 172278880 | 5 | 33951693 | 6 | 50314238 |
| 6 | 76697364 | 6 | 106442646 | 6 | 160701741 | 7 | 4618998 |
| 7 | 12224887 | 7 | 27241878 | 7 | 33247426 | 7 | 51064729 |
| 7 | 69867672 | 7 | 81894122 | 7 | 136662135 | 8 | 57054020 |
| 8 | 105356528 | 8 | 112765092 | 8 | 132586938 | 8 | 145639681 |
| 9 | 81646959 | 9 | 116704056 | 9 | 136769888 | 10 | 95700158 |
| 10 | 106512710 | 11 | 11274951 | 11 | 14477737 | 11 | 27290772 |
| 11 | 28293117 | 11 | 59498158 | 11 | 92211361 | 11 | 106045412 |
| 11 | 121873143 | 12 | 7467944 | 12 | 31095661 | 13 | 35638304 |
| 13 | 60464647 | 13 | 69674812 | 13 | 103529452 | 13 | 105331155 |
| 14 | 35821572 | 14 | 38224432 | 14 | 81267738 | 15 | 72961247 |
| 15 | 28365618 | 16 | 68009705 | 16 | 83022065 | 17 | 1000830 |
| 17 | 50832843 | 18 | 65023688 | 18 | 70830109 | 18 | 77634764 |
| 19 | 6871621 | 19 | 9879584 | 19 | 12082140 | 19 | 51756362 |
| 20 | 26085823 | 20 | 33626399 | 21 | 17552708 | 22 | 21259577 |

## Footnotes

1 https://github.com/CarolaHeinzel/BGA_Classification

2 We employ scikit-learn’s one-vs-rest method to extend ROC AUC to multiclass.

